# High neutralizing potency of swine glyco-humanized polyclonal antibodies against SARS-CoV-2

**DOI:** 10.1101/2020.07.25.217158

**Authors:** Bernard Vanhove, Odile Duvaux, Juliette Rousse, Pierre-Joseph Royer, Gwénaëlle Evanno, Carine Ciron, Elsa Lheriteau, Laurent Vacher, Nadine Gervois, Romain Oger, Yannick Jacques, Sophie Conchon, Apolline Salama, Roberto Duchi, Irina Lagutina, Andrea Perota, Philippe Delahaut, Matthieu Ledure, Melody Paulus, Ray T. So, Chris Ka-Pun Mok, Roberto Bruzzone, Marc Bouillet, Sophie Brouard, Emanuele Cozzi, Cesare Galli, Dominique Blanchard, Jean-Marie Bach, Jean-Paul Soulillou

## Abstract

Perfusion of convalescent plasma (CP) has demonstrated a potential to improve the pneumonia induced by SARS-CoV-2, but procurement and standardization of CP are barriers to its wide usage. Many monoclonal antibodies (mAbs) have been developed but appear insufficient to neutralize SARS-CoV-2 unless two or three of them are being combined. Therefore, heterologous polyclonal antibodies of animal origin, that have been used for decades to fight against infectious agents might represent a highly efficient alternative to the use of CP or mAbs in COVID-19 by targeting multiple antigen epitopes. However, conventional heterologous polyclonal antibodies trigger human natural xenogeneic antibody responses particularly directed against animal-type carbohydrate epitopes, mainly the N-glycolyl form of the neuraminic acid (Neu5Gc) and the Gal α1,3-galactose (αGal), ultimately forming immune complexes and potentially leading to serum sickness or allergy. To circumvent these drawbacks, we engineered animals lacking the genes coding for the cytidine monophosphate-N-acetylneuraminic acid hydroxylase (CMAH) and α1,3-galactosyl-transferase (GGTA1) enzymes to produce glyco-humanized polyclonal antibodies (GH-pAb) lacking Neu5Gc and α-Gal epitopes. We found that pig IgG Fc domains fail to interact with human Fc receptors and thereby should confer the safety advantage to avoiding macrophage dependent exacerbated inflammatory responses, a drawback possibly associated with antibody responses against SARS-CoV-2 or to avoiding a possible antibody-dependent enhancement (ADE). Therefore, we immunized CMAH/GGTA1 double knockout (DKO) pigs with the SARS-CoV-2 spike receptor-binding domain (RBD) to elicit neutralizing antibodies. Animals rapidly developed a hyperimmune response with anti-SARS-CoV-2 end-titers binding dilutions over one to a million and end-titers neutralizing dilutions of 1:10,000. The IgG fraction purified and formulated following clinical Good Manufacturing Practices, named XAV-19, neutralized Spike/angiotensin converting enzyme-2 (ACE-2) interaction at a concentration < 1μg/mL and inhibited infection of human cells by SARS-CoV-2 in cytopathic assays. These data and the accumulating safety advantages of using glyco-humanized swine antibodies in humans warranted clinical assessment of XAV-19 to fight against COVID-19.

## Introduction

Prevention and therapeutic strategies to fight against SARS-CoV-2 comprise, beside steroids and anticlotting agents, vaccination with stabilized messenger RNA, recombinant virus proteins or attenuated virus, the use of antiviral molecules and antibodies targeting the S1 protein. The anti-SARS-Cov-2 antibody passive administration may offer a rapid effect on the virus when administered early in the disease course or complement vaccination by providing an immediate humoral immunity to susceptible persons. Passive immunization with immunoglobulins from convalescent patients is an appealing approach that could be rapidly available provided enough people who have recovered can donate antibody-containing serum. Experience from prior outbreaks with other coronaviruses, such as SARS-CoV, shows that passive immunization with convalescent plasma does control viremia and confers clinical benefit ^1^. Passive antibody therapy has been invented in the 19^th^ century when it was the only means of treating certain infectious diseases prior to the development of antibiotics ^2^. Interestingly, passive immunotherapy with heterologous animal sera had been concomitantly used for treating solid tumors and clinical responses had also been reported ^3^. Today, several passive heterologous immunotherapy products including polyclonal and monoclonal antibodies against infectious agents are on the market and have demonstrated robustness and efficacy to fight against bacterial infections, including tetanus, botulism, diphtheria, or viral infections such as hepatitis A and B, varicella ^4^, rabies ^5^. Others are in development ^6,7^. CP from COVID-19 patients are currently tested worldwide ^8^. The anticipated mechanism of action by which antibodies would mediate protection against viruses is neutralization, i.e. inhibition of viral particles interaction with their cognate receptor and, thereby, entry into human cells. However, other anti-viral mechanisms involving antibodies may be possible, such as antibody-dependent cellular cytotoxicity and/or phagocytosis. In the context of enveloped viruses, such as coronaviruses, direct complement-mediated lysis might also participate in viral particles elimination since expression by viral particles of complement-regulatory proteins belong to the viral-mediated immune escape mechanisms ^9^.

Whereas polyclonal antibodies can present potent neutralizing effect on receptor binding and direct lysis properties against viral particles ^9^, administration of heterologous animal-derived immunoglobulins may elicit serious adverse effects in humans such as serum sickness disease (SSD) almost always present in immunocompetent hosts ^10^ and may provide “xenosialitis”, a systemic inflammation still elusive in human but associated with cardiovascular diseases and cancer in animal models ^11,12,13^.

The description of serum sickness more than a century ago in humans perfused with animal sera ^14^ eventually led to the identification of a class of human antibodies directed against glycans bearing the common mammalian sialic acid N-Glycolylneuraminic acid (Neu5Gc) ^15,16^. Humans are with few exceptions the only known mammals presenting a biased terminal sialylation of glycoproteins and glycolipids. They lack Neu5Gc on glycosylated proteins or lipids, due to a human-lineage specific genetic mutation in the enzyme cytidine monophosphate-N-acetylneuraminic acid hydroxylase (CMAH). A direct consequence is that Neu5Gc epitopes are excluded from « self-tolerance » and natural anti-Neu5Gc antibodies are present. Furthermore, new antibodies can be easily elicited following infusion of Neu5Gc-positive proteins of animal origin, including therapeutic immunoglobulins. Humans also lack the α1,3-galactosyl-transferase enzyme (GGTA1), are not tolerant to α1,3-galactose (αGal) epitopes, present various levels of natural anti-αGal antibodies and increase their level of anti-αGal antibodies after infusion of animal-derived products ^17^. Rabbit anti-lymphocyte serum treatment in diabetic patients, for example, resulted in highly significant increases of both anti-αGal and anti-Neu5Gc IgM and IgG, peaking at 1 month and still detectable one-year post infusion, that were responsible for the induction of SSD, an immune-complex disease, in almost all patients^18^.

Initially developed for xenotransplantation purposes, pigs have been genetically engineered to knock out the CMAH and GGTA1 enzymes. Immunoglobulins from these animals are devoid of Neu5Gc and αGal carbohydrate epitopes and may not elicit SSD and possible deleterious anti-xenoantigen antibodies when administered in human. The limited clinical experience gained so far with such glyco-humanized swine polyclonals, lacking both Neu5Gc and αGal sugar moieties, used to confer immunosuppression in kidney transplant recipients suggests their good tolerance and efficacy (NCT 04431219).

Another feature of such swine glyco-humanized polyclonals revealed here is their inability to bind human FcγR (CD16, CD32, CD64) which should protect recipients from antibody-dependent enhancement (ADE), a process by which several viruses (among which dengue, zika, SARS-CoV) have an increased capacity to enter into cells expressing FcγR (mainly macrophages) in the presence of virus-specific antibodies or cross-reactive antibodies ^19,20^. However, whether infection with SARS-CoV-2 might elicit ADE remains speculative in human ^21^. Failure to bind human FcγR should also limit FcγR-dependent recruitment/activation of macrophages described to contribute in skewing inflammation-resolving response during SARS-CoV infection ^22^. Here, we described the immunization of CMAH/GGTA1 DKO pigs with a recombinant RBD domain from SARS-CoV-2 spike protein and the development of XAV-19, a drug candidate manufactured from hyperimmune serum presenting a higher therapeutic index than convalescent plasma.

## Methods

### Animals, reagents

Pigs double knocked out for GGTA1 and CMAH encoding genes ^23^ were obtained by cloning procedures described in Lagutina et al 2006 ^24^. All procedures involving the use of animals in this study were carried out in accordance with the Italian Law (D.Lgs 26/2014) and EU directive 2010/63/EU regulating animal experimentation after authorization by relevant authorities (Italy, project n 959/2017☐PR; Belgium 2010/63/EU and AR 29/05/2013). The invalidating mutations in genes encoding the CMAH and GGTA1 were confirmed by direct amplification and sequencing of the gene sequences at the animal facility, to ensure the stability of the genetic background. The phenotype has been checked by mass spectrometry to confirm the absence of αGal and Neu5Gc residues on gamma immunoglobulins. Recombinant proteins from SARS-CoV-2 have been manufactured from Hek-293 cells and were purchased from Interchim, Monluçon, France.

### LC-MS analysis of glycosylation

Measurement of N-glycan Neu5Gc and αGal on DKO and WT porcine IgGs has been performed by deglycosylation, N-glycan procainamide labeling protocol and treatment with α-galactosidase, were as previously described ^25^.

### Analysis of human anti-pig IgG reactivity

Pig WT IgG, IgG from GGTA1 KO, IgG from CMAH/GGTA1 double knock out (DKO) pigs or rabbit IgG (WT) have been coated on Maxisorp™ (NUNC) ELISA plates at 10 μg/mL overnight at 4°C in carbonate buffer pH 9.5. After washing with PBS-Tween 20 (0.05%), free binding sites were blocked with 1% chicken egg albumin. Anonymously coded human serum samples from healthy volunteers (from the Etablissement Français du Sang, Nantes, France) were incubated at a 1:100 dilution, for 2h at 37°C. After washing, bound IgG molecules were revealed with peroxidase conjugated goat anti-human IgG antibodies (Jackson ImmunoResearch, 1/10 000 in 1% chicken egg albumin) and TMB reagent. Optical density was read at 450 nm.

### Complement-mediated cytotoxicity

Rabbit anti-T cell globulins (Thymoglobulin^®^) and glyco-humanized pig anti-T cell globulins were differentially diluted in non-immune IgG solutions to obtain preparations presenting similar binding titers to human T cells. Then, serial dilutions were added to 250 000 human PBMC at 4°C for 30 minutes. After washing, human serum used as a source of complement diluted 3 times in cold RPMI medium was added, on ice, on each well. Reaction was then initiated by placing tubes at 37°C, and maintained for 30 min. After this step, the pellets were placed on ice and resuspended with propidium iodide, which is a viability marker of cells and the cytotoxic activity of antibodies was determined by flow cytometry, after gating on lymphocytes, based on their morphology.

### Binding to human FcγR

The binding of IgG from DKO swine to the FcγRI, FcγRIIa and FcγRIIb human receptors was tested using an ELISA assay and BIAcore surface plasmon resonance (SPR). IgG from different species were purified by protein A affinity and coated on Maxisorp™ ELISA plates overnight at 4°C (100μL/well, 20μg/mL in carbonate buffer pH 9.5). After washing with PBS-Tween 20 0.05%, free binding sites were saturated with 2% bovine serum albumin. His-Tagged FcγRI, FcγRIIa and FcγRIIb receptors (CD64/32a/32b, R&D Systems; 100μL, 2.4 μg/mL in 2% bovine serum albumin) were then added and incubated for 2h at room temperature (RT). After washing, bound FcγR molecules were revealed with peroxidase conjugated anti-tag antibody (Miltenyi, 120-003-811; 1/5000 in 2% bovine serum albumin) and tetramethylbenzidine (TMB) reagent. Optical density was read at 450 nm and 630 nm. Specific optical density is the value read at 450 nm minus that read at 630 nm. The binding of pig and rabbit (Thymoglobulin®, Sanofi) IgG to human FcγRIIIa (CD16a, R&D Systems, 4325-FC) receptor was also measured by SPR. IgG were immobilized on a CM5 sensor chip by amine coupling to a level of 2000RU. Free reactive sites were then inactivated with ethanolamine 1M pH 8.5. Different concentrations (5μM, 2.5μM, 1.25μM, 0.625μM, 0.312μM, 0.156μM) of human FcγRIIIa were injected on the chip, with association time intervals of 180s and dissociation time of 600s. Regeneration between cycles were performed by 100mM NaOH treatment for 45s.

### Anti-SARS-CoV-2 binding ELISA

The target antigen (SARS-CoV-2 Spike RBD protein, Interchim) was immobilized on Maxisorp plates at 1μg/mL in carbonate/bicarbonate buffer at 4°C overnight). After washing, saturation was performed with PBS-Tween-BSA for 2h at room temperature. Samples were diluted into PBS-Tween and added into the plate in duplicate, incubated 2h at RT and washed 3 times. Bound pig IgG were revealed with a secondary anti-pig-HRP-conjugated antibody (Bethyl Laboratories, USA) diluted in washing buffer, at 1:1000, incubated 1h at RT and washed 3 times. TMB reagent was added in the plate, incubated up to 20 minutes in the dark and stopped with H_2_SO_4_. Reading was performed at 450nm.

### Inhibition of Spike/ACE-2 interaction

An assay was developed to assess the properties of swine antibodies to inhibit binding of a recombinant SARS-CoV-2 spike S1 molecule to immobilized recombinant ACE-2 molecules. The target antigen (human ACE-2, Interchim) was immobilized on Maxisorp plates at 1 μg/mL in carbonate/bicarbonate buffer at 4°C overnight). The plates were washed in PBS-Tween-0.05% and saturated with PBS-Tween-0.05%-2% skimmed milk for 2h at RT. Ligand Spike S1-hFc tag (Interchim) at 125 ng/mL in PBS-Tween-0.05%-1% skimmed milk was then added into another plate in duplicate, and pre-incubated 30min at RT in the presence of test samples diluted in PBS-Tween-0.05%-1% skimmed milk powder (range between 50 and 0.39 μg/mL). Then the mixture was added to the plate for a 2h incubation at RT. The human Fc tag was then revealed with a specific HRP-conjugated anti-human IgG secondary antibody (diluted in in PBS-Tween-0.05%-1% skimmed milk powder at 1:1000, incubated 1h at RT and washed 3 times). TMB reagent was added in the plate, incubated up to 30 minutes in the dark and stopped with H_2_SO_4_. The plate was read at 450 nm.

### Plaque Reduction Neutralization Testing (PRNT)

The plaque-reduction neutralization test (PRNT) was performed in duplicate using 24-well tissue culture plates (TPP Techno Plastic Products AG, Trasadingen, Switzerland) in a biosafety level 3 facility. Swine serum dilutions were mixed with equal volumes of SARS-CoV-2 (BetaCoV/Hong Kong/VM20001061/2020 [KH1], corresponding to the D614 variant) or SARS-CoV (strain HK39849, SCoV) at a dose of 200 plaque-forming units determined by Vero E6 cells. Briefly, after 1 hour of incubation at 37°C, the virus-serum mixture was added to Vero E6 cell monolayers in 24-well tissue culture plates. After 1 hour of adsorption, the virus-serum mixture was removed, and plates were overlaid with 1% agarose with cell culture medium. Plates were then incubated for 3 days at 37°C in 5% CO_2_ in a humidified incubator before fixation with 4% paraformaldehyde and staining crystal violet solution. The highest plasma dilution that resulted in 90% reduction of plaque numbers was denoted as PRNT90. A virus back-titration of the input virus was included in each batch of tests.

### Cytopathogenic Effect (CPE) assay

To further analyze the neutralization potency and to confirm data with another independent virus strain, a CPE assay was carried out at VibioSphen, Labège, France. The purpose of this study was to assess the ability of the serum pool, XAV-19 R&D, and XAV-19 batches BMG170-B01 to BMG170-B04 to inhibit entry and cytopathic impact of SARS-CoV-2 on sensitive cells (Vero E6 Cells). All experimental protocols involving live SARS-CoV-2 followed the approved standard operating procedures of the Biosafety Level 3 facility. SARS-CoV-2 was isolated from a patient with laboratory-confirmed COVID-19 at VibioSphen, corresponding to the G614 variant. The viral isolate was amplified by one additional passage in VeroE6 cells to make working stocks of the virus. Vero E6 Cells were cultured in Dulbecco’s modified Eagle’s medium (DMEM) supplemented with 10%v/v fetal bovine serum, 1%v/v penicillin-streptomycin supplemented with 1%v/v sodium pyruvate at 1x 105 cells per well in 12-well tissue culture plates. At 100% confluence (2 days post-seeding), the cells were washed twice with PBS and six serial dilutions of the virus (1/10 each time) will added to the cells. Following infection with 0.3 ml per well of each dilution, plates were incubated at 37°C for 1 h, and the cells were washed with PBS before the addition of 2% w/v agar containing 1 μg/mL-5 tosyl phenylalanyl chloromethyl ketone-trypsin (Sigma-Aldrich,) to the cell surface. Plates were left at RT for 20–30 min to allow for the overlay to set, and were then incubated at 37°C for 72 h. Cells were fixed with 4% v/v paraformaldehyde before both the fixative and agar will be removed and the cells stained with 0.1%w/v Crystal Violet (Fisher) in 20%v/v ethanol. Plaque titers were determined as plaque forming units per mL.

CPE reduction assay was performed as follows: Vero E6 cells were seeded in 96-well clusters at a density of 5000 cells/well 2 day before infection. Two☐fold serial dilutions, starting from 200 or 250 μg/mL or from a 1/25 dilution of the serum were then mixed with an equal volume of viral solution containing 100 TCID50 of SARS☐CoV☐2. The serum☐virus mixture was incubated for 1 hour at 37°C in a humidified atmosphere with 5% CO2. After incubation, 100 μL of the mixture at each dilution was added in triplicate to a cell plate containing a semi☐confluent VERO E6 monolayer. The plates were then incubated for 3 days at 37°C in a humidified atmosphere with 5% CO_2_. Cell controls were infected with SARS-CoV-2 at a multiplicity of infection of 0.01. The incubation period was 72h. Uninfected cells were included as controls to exclude potential cytotoxic/cytostatic effects of the compound treatment. All samples were tested in duplicate.

### Statistical analysis

Continuous variables are expressed as mean ± SEM, unless otherwise indicated, and compared with the nonparametric Mann-Whitney two-sided test or Kruskal–Wallis tests with Dunn’s ad hoc pairwise comparisons for more than two groups. P values of <0.05 were considered to be statistically significant. All statistical analyses were performed on GraphPad Software (GraphPad Software, San Diego, CA).

## Results

### Absence of αGal and Neu5Gc carbohydrate on immunoglobulins from CMAH/GGTA1 double knockout swine selected to elicit an immune response against SARS-CoV-2 spike RBD domain

We have already shown that IgG from CMAH and GGTA1 DKO pigs were deprived of Neu5GC and αGal carbohydrates ^26^. To confirm the absence of these carbohydrates on IgG from animals immunized with the SARS-CoV-2 spike RBD domain to produce XAV-19, the clinical batch of anti-CoV IgG, N-glycan profiling was carried out as previously described ^27,26,23^, and compared with wild-type pig IgG. The absence of N-glycan structure exhibiting Neu5Gc was clearly confirmed according to two independent methods: N-glycan profiling by ultraperformance liquid chromatography and ESI-MS detection (UPLC-FLD/ESI MS) after enzymatic deglycosylation by PNGase F on one hand, and quantitative analysis of sialic acid by HPLC after DMB labeling on the other hand. Instead, the human-type acetylated form of neuraminic acid Neu5Ac was observed (Figure 1). Limited by the sensitivity of the method, no N-glycan with αGal moieties was observed and detected on glyco-humanized pig IgG (data not shown).

**Figure 1:**
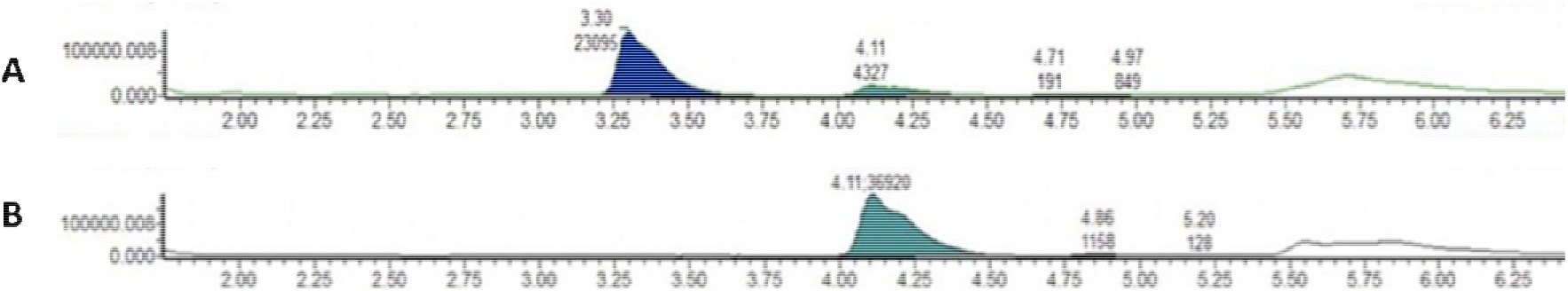
RP-UPLC chromatograms of pig IgG and the clinical batch of glyco-humanized pig IgG showing absence of Neu5Gc residues on glyco-humanized pig IgG. Sialic acids quantification was performed by DMB-labeled released sialic acid analysis by RP UPLC-FLD/ESI-MS. Labeled monosaccharides were separated by UPLC using a BEH C18 column and the quantification was performed with a fluorescent detector coupled to the chromatography. The procedure was performed following standard procedures from Waters (Application Note UPLC/FLR/QT of MS Analysis of Procainamide-Labeled N-glycans).Wild type pig IgG show Neu5GC and Neu5AC peaks (panel A) whereas glyco-humanized pig IgG only shows a Neu5AC peak (panel B). Filled histograms denote the relative amount of each N-glycan in the sample.

### Reduced recognition of glyco-humanized swine IgG by human natural antibodies

Formation of immune complexes between pre-existing or elicited anti-Neu5Gc antibodies and therapeutic immunoglobulins of animal origin has been identified as a major trigger of serum sickness after serotherapy ^15,16,28,18,29^. To explore the actual impact of removal of αGal and Neu5Gc epitopes on recognition of swine IgG by pre-existing human natural antibodies, human serums from healthy volunteers were assessed in a binding assay against wild-type or glyco-humanized swine IgG, or against rabbit IgG used as a comparator. The data revealed an overall lower reactivity of human natural antibodies against pig IgG than against rabbit IgG. Reactivity against glyco-humanized swine IgG was considerably reduced compared with reactivity against wild-type IgG (p<0.001). Interestingly, this difference was due to the lack of Neu5Gc and not to the lack of αGal, since recognition of IgG from WT or GGTA1 single KO animals was similar (Figure 2), fitting to the previously reported ^26^ low αGal loading of pig IgG on mass spectrometry pattern. The importance of the two terminal sugar epitopes in elicited responses against foreign IgG has been also well documented elsewhere ^18,28,29^.

**Figure 2:**
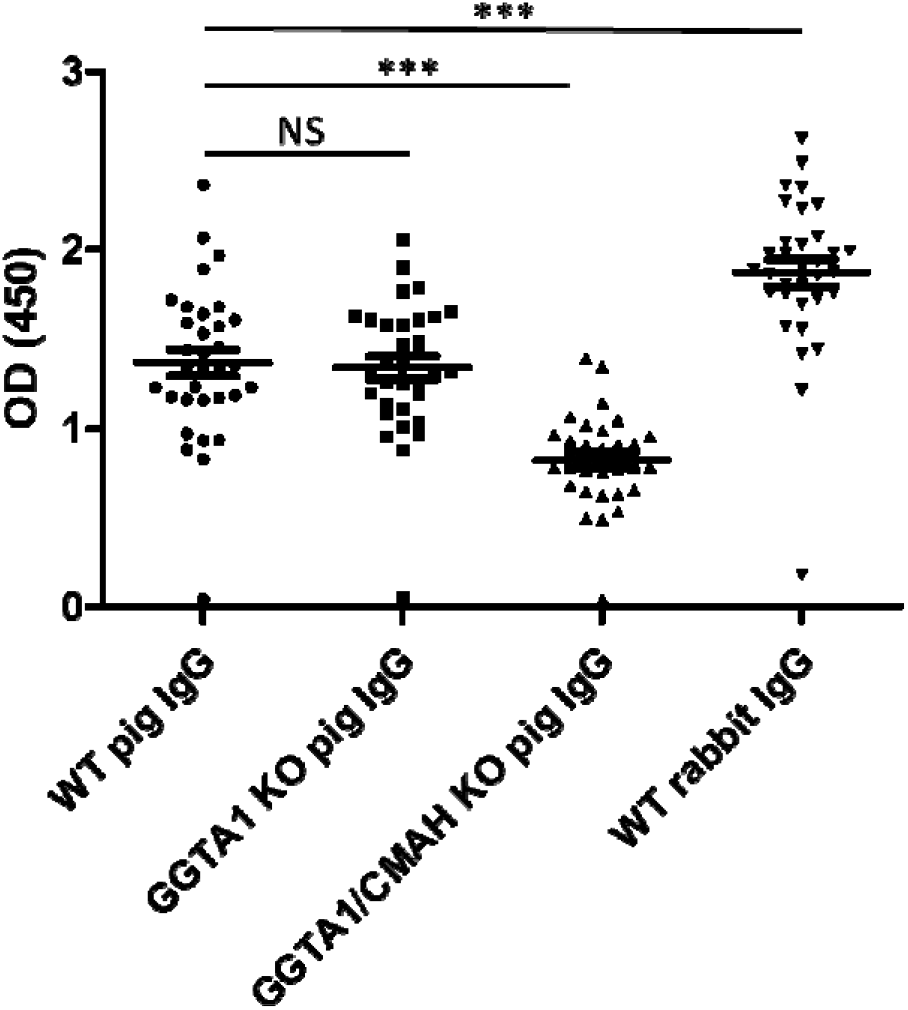
Reactivity of human IgG against rabbit, pig and CMAH/GGTA1 DKO pig IgG. Serum from healthy volunteers (n=59; 100-fold dilutions) was incubated on ELISA plates coated with wild-type pig IgG, GGTA1 KO pig IgG, CMAH/GGTA1 DKO pig IgG or rabbit IgG. Binding of human IgG was revealed with a labeled secondary antibody against human IgG. NS: not significant. ***, p<0.001.

### IgG from CMAH/GGTA1 DKO pigs show high complement activation capacity but no binding to human FcγR

We observed that the IgG fraction of CMAH/GGTA1 DKO pigs presented an overall higher complement-activating capacity when compared with rabbit immunoglobulins. This could be evidenced by immunizing CMAH/GGTA1 DKO pigs with human T cells and assessing complement-mediated cytotoxicity (CDC) of the IgG fraction, in comparison with a rabbit IgG anti-T cell immunoglobulin (Thymoglobulin^®^). We used adjusted concentration of pig and rabbit IgG preparations to obtain identical binding intensity to target cells by flow cytometry (Figure 3A). The data showed a CDC activity of Neu5Gc/αGal-negative pig IgG higher than with rabbit IgG (Figure 3B). To further investigate whether Neu5Gc/αGal-negative pig IgG inherently activate more complement than rabbit IgG, independently of their antigen target recognition capacities, we directly assessed complement component 1q (C1q) binding intensity to immobilized IgG. This experiment confirmed higher recruitment of complement C1q by Neu5Gc/αGal-negative pig IgG than by rabbit IgG (Supplementary Figure 1A, B). To understand whether this high complement-activation activity was a consequence of the altered glycosylation of Neu5Gc/αGal-negative pig IgG (glycosylation being required for efficient C1 activation ^30^), we also compared C1q binding to immobilized WT and Neu5Gc/αGal-negative pig IgG. The data revealed similar C1q recruitment (Supplementary Figure 1C, D), suggesting absence of intrinsic difference due to conversion of Neu5Gc to Neu5Ac type of N-glycosylation.

**Figure 3:**
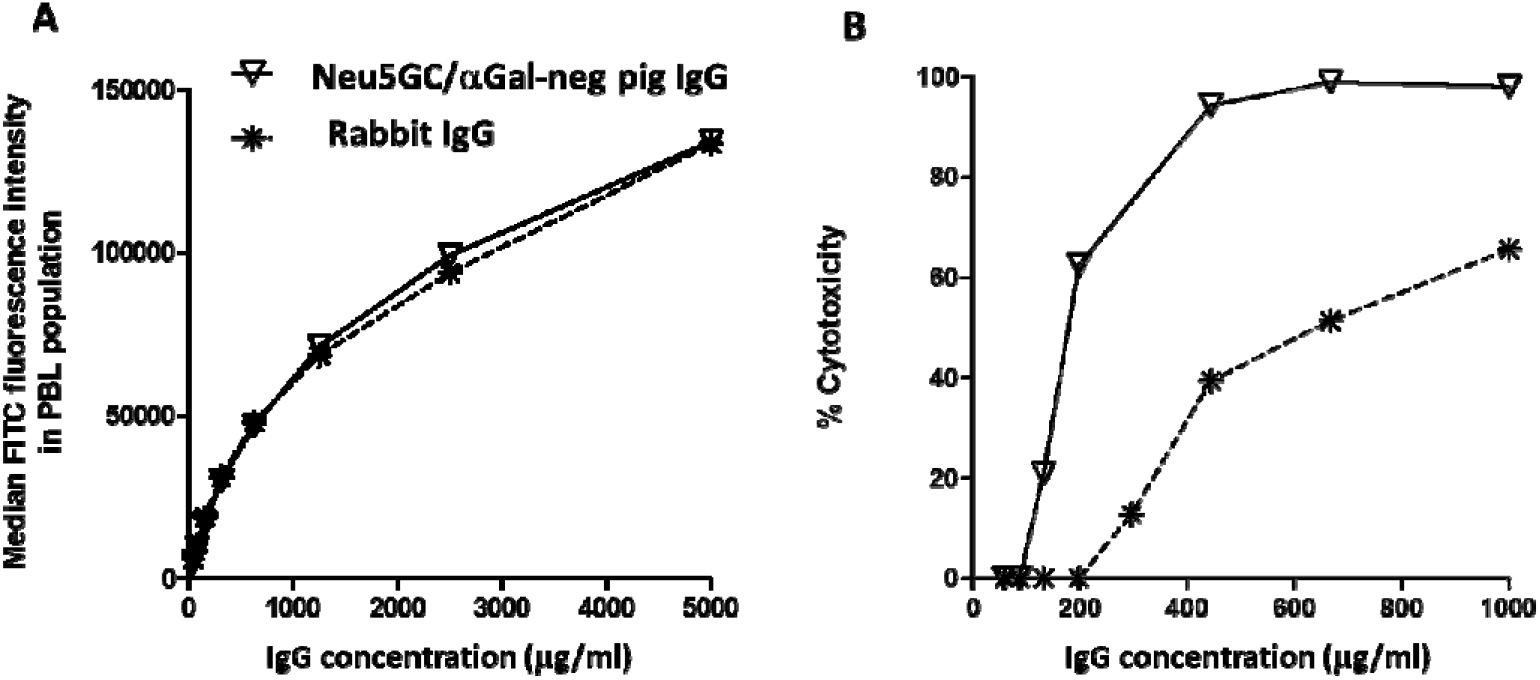
High complement activation with Neu5Gc and αGal-negative pig IgG. GGTA1/CMAH double KO pigs were immunized with human T cells and the hyperimmune fraction purified by Protein A chromatography. Rabbit anti-human T cells used here was Thymoglobulin®. The two IgG preparations were differentially diluted in non-immune IgG of the respective species to obtain similar binding titer and intensity on target human T cells by flow cytometry. A fluorescent Protein-G reagent has been used instead of species-specific secondary antibodies to ensure similar, reagent-independent revelation (A). Target human T cells were used in a Complement-mediated cytotoxicity assay where rabbit IgG and GGTA1/CMAH double KO pig IgG directed against human T cells were compared (B). Results shown here are representative of 3 independent experiments.

In parallel, using either wild or the DKO pig-derived IgG preparations against human T cells, we noticed the absence of antibody-dependent cell-mediated cytotoxicity (ADCC) using human effector cells, when compared with rabbit IgG directed against the same target cells (Supplementary Figure 2). This led us to investigate interaction between DKO pig IgG and human FcγR (CD16, CD32 and CD64). Surface plasmon resonance and ELISA experiments revealed absence of interaction of pig IgG, either DKO or wild-type, with human CD16, CD32 or CD64, whereas binding of IgG from the rabbit, cow, horse, donkey, goat and human all presented reactivity against at least one Fc receptor (Table 1 and Supplementary Figure 3).

**Table 1:** Interaction of polyclonal IgG from the indicated species with human FcγR.

| | FcγRIII (CD16) | FcγRIIa (CD32a) | FcγRIIb (CD32b) | FcγRI (CD64) |
| --- | --- | --- | --- | --- |
| pig | - | - | - | - |
| GH-pig | - | - | - | - |
| rabbit | + | + | + | + |
| bovine | ND | + | + | + |
| horse | ND | - | - | + |
| donkey | ND | - | - | + |
| goat | ND | + | + | + |
| human | + | + | + | + |

ND: not done

### Anti-SARS-CoV-2 hyperimmune serum titers

CMAH/GGTA1 KO pigs were immunized by intramuscular (IM) administrations of SARS-CoV-2 RBD spike antigens in mineral adjuvant. The RBD sequence was selected based on recent demonstration that it is instrumental in the binding of the spike protein to ACE-2 ^31^ and that antibodies to RBD consequently inhibit SARS-CoV-2 entry into ACE-2-positive cells ^32,33^. Immune serum presented shortly after immunization end binding titers above 1:40,000 (Figure 4A). After several immunizations, end titers in individual animals ranged between 1:100,000 and >1:10^6^ (Figure 4B). IgG extracted from sera by Protein A chromatography presented an initial EC50 in ELISA of 2.5 μg/mL after two immunization and below 1 μg/mL after 3 or more immunizations (Figure 4C, D).

**Figure 4:**
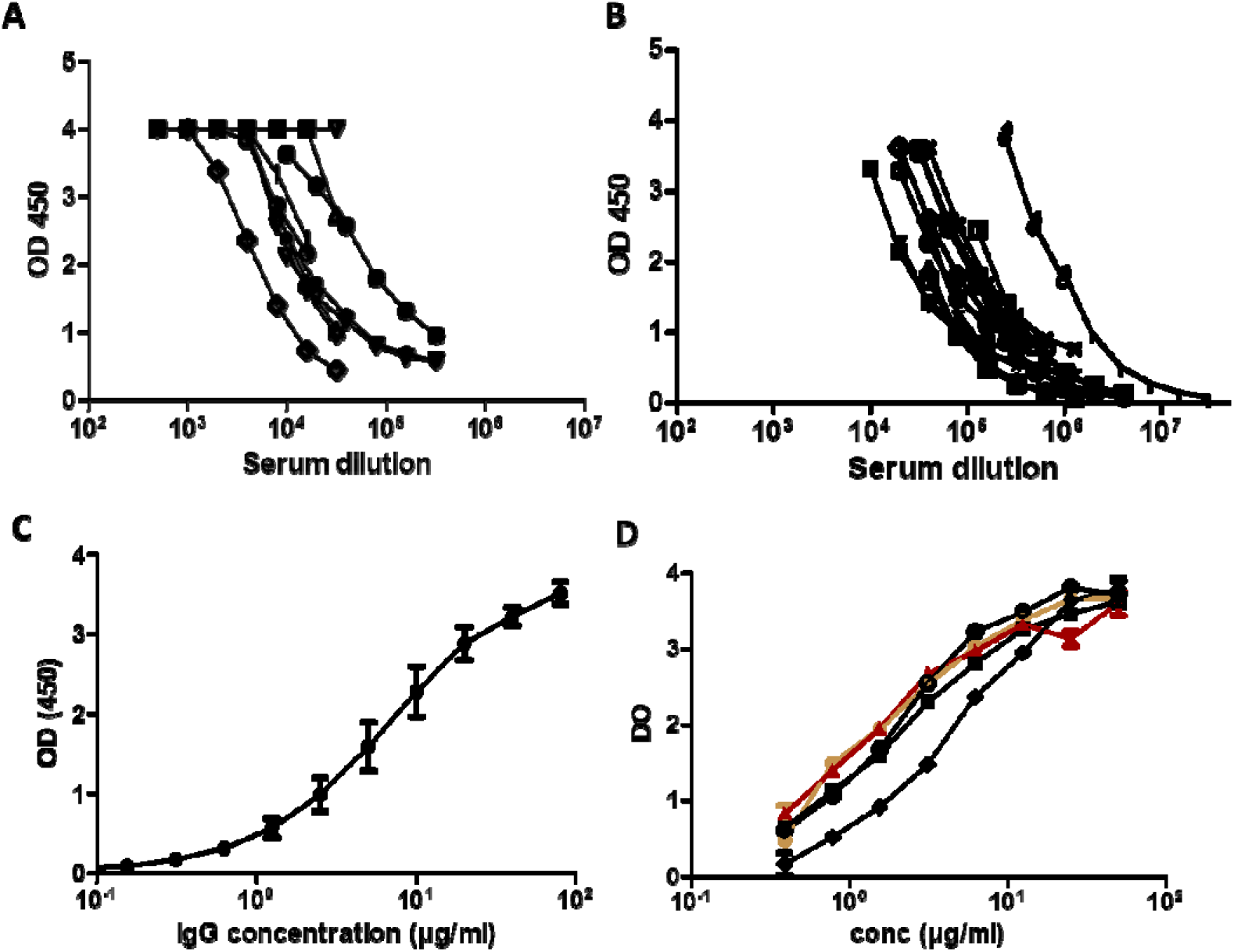
SARS-CoV-2 spike binding ELISA. GGTA1/CMAH double KO pigs were immunized with SARS-CoV-2 spike RBD recombinant proteins and blood was collected after 2 (A, C) or 3 to 5 (B, D) immunizations. Serum (A, B) or the IgG fraction (C, D) were tested by ELISA for binding to recombinant SARS-CoV-2 spike molecules. In A, B and D, curves represent data from individual animals. In C, IgG were extracted from a serum pool from animals depicted in A (Means ±SD of triplicates). In D, the orange curve corresponds to one of the best responding animals and the red one to the pool of sera selected for further GMP processing and clinical usage.

### Inhibition of Spike/ACE-2 interaction

Anti-SARS-CoV-2 hyperimmune sera were also tested in a spike/ACE-2 binding competition assay. Hyperimmune sera demonstrated an inhibitory capacity with an end titer of 1:4000 soon after immunization (Figure 5A) and IgG extracted from sera by Protein A chromatography presented an IC50 of 2.5 μg/ml (Figure 5B). After a few repeated immunizations, serum presented inhibitory capacities at much higher dilutions and IgG from these sera could be diluted down to < 0.1 μg/mL (Figure 5C, D).

**Figure 5:**
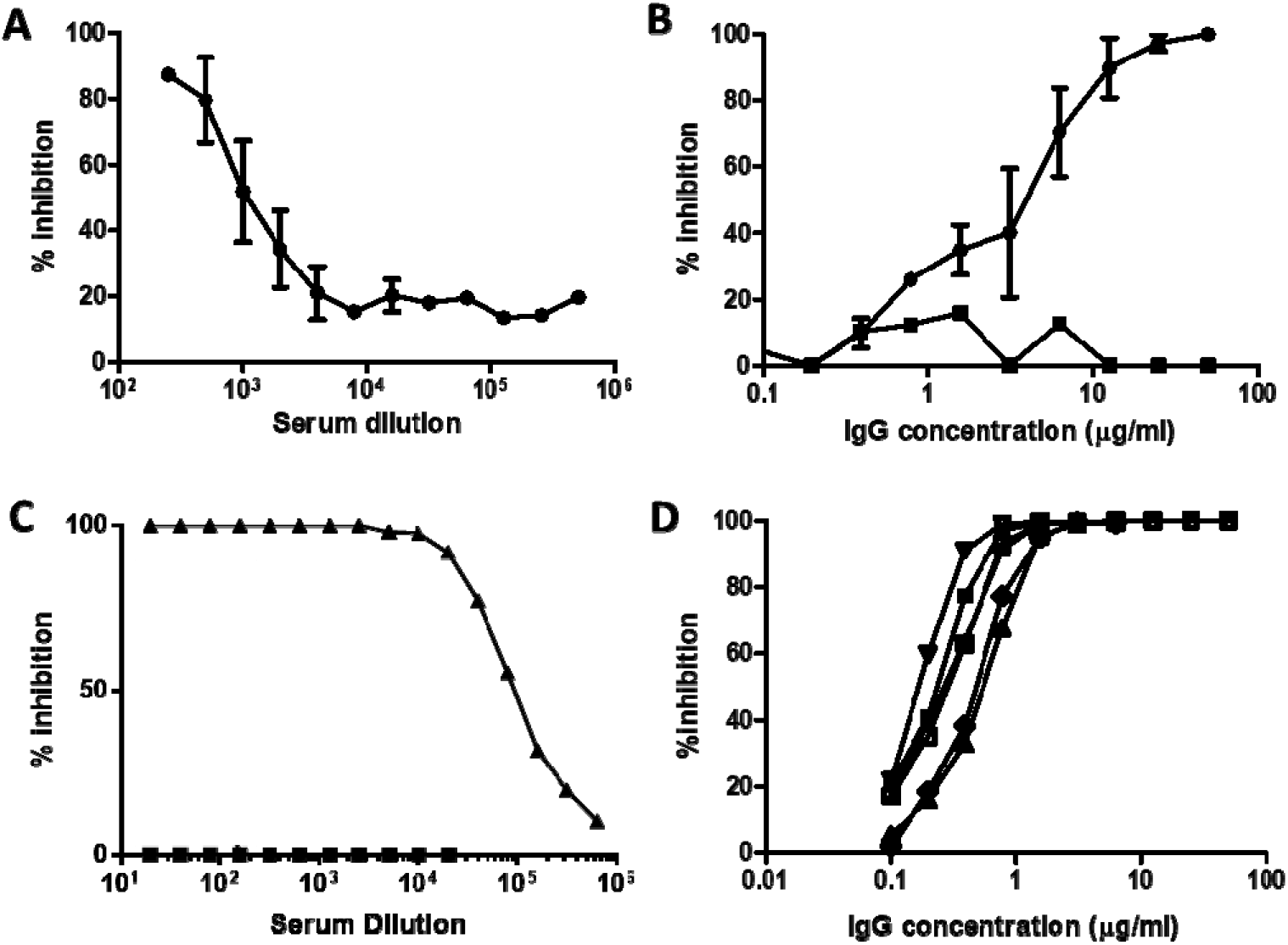
Assessment of inhibition of SARS-CoV-2 Spike/ACE-2 interaction. Samples analysed in Figure 4 were tested in an ELISA interaction assay where ACE-2 was immobilized on plastic and Spike-Fc ligand binding to ACE-2 was revealed with a secondary antibody against Fc. 100% inhibition represents absence of Spike/ACE-2 interaction. A: Mean ± SD of 4 individual sera collected after 2 immunizations. B: Mean ± SD of individual IgG fractions shown in A. Black square symbols: IgG extracted from a pool of pre-immune serum. C: Triangles represent data from a serum pool obtained after 3 to 5 immunizations. Square symbols: non-immune serum. D: Mean ± SD of individual IgG fractions shown in C.

### Neutralization of SARS-CoV-2 by glyco-humanized swine IgG

The neutralizing effect of the hyperimmune serum and IgG fractions against live SARS-CoV-2 virus (G614 form) was determined by CPE reduction assay. CPE_100_ (maximal dilution to reach 100% of neutralization) with pooled hyperimmune sera used to prepare therapeutic IgG reached 1:1600 and the concentration of the IgG preparation required to reach CPE_100_ ranged between 15 and 25 μg/mL for the samples collected after 2 immunizations. CPE_100_ was reached at 3.125 μg/mL when using IgG preparations collected after 3 immunizations or more (Table 2).

**Table 2:** Neutralizing activity by Cytopathogenic Effect (CPE) assay and Plaque Reduction Neutralizing Test (PRNT).

|  | **Serum pool\*** | **Purified IgG from serum pool** |
| --- | --- | --- |
| End titer or concentration (µg/ml) to abrogate CPE (100% efficacy) against SARS-CoV-2 – samples collected after 2 immunizations | 1:1600 | 15 to 25 µg/ml |
| Concentration (µg/ml) to abrogate CPE (100% efficacy) against SARS-CoV-2 – samples collected after 3 immunizations or more | ND | 3.125 µg/ml |
| End titer or concentration (µg/ml) to abrogate PRNT (90% efficacy) against SARS-CoV-2 – samples collected after 2 immunizations | >1:320 | ND |
| End titer or concentration (µg/ml) to abrogate PRNT (90% efficacy) against SARS-CoV – samples collected after 2 immunizations | 1:10** | ND |

\* corresponding to the sample tested in Figure 5A
\*\* another sample drawn after several immunizations presented a PRNT<sub>90</sub> at the dilution of 1:80 against SARS-CoV
ND: not done

The neutralizing effect of the hyperimmune serum was also determined in parallel by plaque reduction neutralization test (PRNT) using another SARS-CoV-2 strain (presenting the D614 form), as previously described ^34^. Serum samples collected from three pigs immunized twice showed PRNT_90_ at 1:160, 1:80 and >1:320 against SARS-CoV-2, the serum collected from the pig displaying the highest titer by PRNT90 (>1:320) also cross-neutralized SARS-CoV at PRNT90 1:10. The first pool of serum obtained after repeated immunizations, from which IgG have been extracted to prepare the initial clinical batch of neutralizing immunoglobulins, presented a PRNT_90_ titer of >1:320 and cross-neutralized SARS-CoV at PRNT_90_ 1:80 (Table 2).

## Discussion

The ongoing COVID-19 pandemic requires prompt availability of prophylactic and therapeutic interventions. Preliminary data on transfer of convalescent plasma (CP) from COVID-19 recovering patients to critically ill patients demonstrated efficacy to reduce viral titers and to improve clinical symptoms when provided early in the disease course ^35,36^. The effect has been attributed to the presence of neutralizing antibodies against the virus spike RBD sequence, since antibodies uniquely directed against RBD do protect mice in preclinical studies ^37^. Yet CP show highly variable neutralization half-titers of anti-SARS-CoV-2 antibodies, ranging from less than 1:50 in a third of patients, between 1:50 and 1:1,000 in the majority of them, with only rare patients showing titers above 1:5,000 ^38^.

In the past clinical experience with CP from SARS-convalescent patients in 2003 (80 patients), the average CP volume was 279 mL and the neutralizing titer was 1:160 ^39^. Today, in critically ill patients with COVID-19, Duan et al ^40^ reported using one dose of 200 ml of CP presenting a neutralizing titer above 1:160. Shen et al ^41^ reported efficacy after two consecutive transfusions with 200 mL of CP with end point dilution titers greater than 1:40, corresponding to a SARS-CoV-2–specific antibody (IgG) binding titer greater than 1:1000. In this report, the ratio between binding and neutralization titers is therefore of 25-fold on average. Recent guidelines from the European Commission on the use of CP revised the desired neutralizing CP titers and suggested that “neutralizing antibody titers should optimally be greater than 1:320, but lower thresholds might also be effective” (https://ec.europa.eu/health/blood_tissues_organs/covid-19_en). Thus, altogether, estimating that transfusion of 200 mL CP corresponds to the administration of approximately 25 mg/kg IgG (considering that plasma contains 10 mg/mL IgG and considering 80 kg as an average patient’s weight), available data suggest that 25 mg/kg of an IgG preparation with a neutralizing titer of at least 1:40 meets the clinically required efficacy.

A few weeks after immunization of CMAH/GGTA1 KO pigs with SARS-CoV-2 RBD proteins, binding and neutralization serum antibody titers rapidly reached 1:100,000 and 1:4,000 (end titer dilutions in the ELISA neutralization assay), respectively, corresponding to the same 25-fold ratio described in humans. The neutralization titer in a cytopathogenic effect assay was of 1:1600 but corresponds to the dilution required to reach 100% inhibition (as opposed to end-titer dilution). The neutralizing capacity was confirmed in a PRNT_90_ assay with another, independent SARS-Cov-2 strain (showing PRNT_90_ > 1:320). This neutralizing titer is at least 40-fold higher (depending on the assay being considered) as compared to COVID-19 CP presenting a clinical interest according to the literature, and at least 12.5-fold higher than the European Commission recommended. IgG purified from pig hyperimmune serum could have therefore the potential to be used successfully at doses 12.5 to 40-fold lower than CP, i.e. at doses starting at 0.6 mg/kg.

Glyco-humanization refers to removal of the αGal epitopes and conversion of animal-type Neu5Gc/Ac forms of sialic acid into human single type Neu5Ac. The procedure has been initially developed to circumvent problems related to the occurrence of high titers of anti-αGal and anti-Neu5Gc antibodies following administration of anti-T lymphocytes rabbit immunoglobulins, which is potentially causing allergies ^42^, serum sickness ^28^ and a possible “xenosialitis”, a systemic inflammation so far only demonstrated in animal models ^11,12^. Although our experiments do not directly address the clinical importance of avoiding anti-αGal/Neu5Gc antibody responses, our data show that “natural” anti-pig antibodies found in human serum are essentially directed against Neu5Gc and that reactivity against rabbit IgG is higher than against pig IgG (Figure 2). These data are of importance in a context where hyperimmune pig immunoglobulins are being used therapeutically to treat aplastic anemia ^42,43^ and are being developed as an anti-lymphocyte induction treatment in kidney transplantation (NCT 04431219). Glyco-humanized pig immunoglobulins would potentially present, therefore, a safety advantage over rabbit polyclonal antibodies or over immunoglobulins from other species currently in clinical use, such as bovine, horse and goat. Another potential advantage of using glyco-humanized antibodies from pigs is related to its unique mechanism of action in humans. Because these modified pig IgG fail to interact with human Fc receptors and to recruit human effectors (Table 1 and Supplementary Figure 2 and 3), likely due to divergent molecular evolution, they are not susceptible to promote inflammatory-related ill-effects associated with IgG Fc/ FcγR interactions. One such interaction demonstrated with diverse viral families, including Coronaviridae is ADE ^44,45,46^. However, the importance of ADE phenomenon, whereby a previous immune response could render an individual more susceptible to a Cov-2 reinfection, remains undetermined ^47,21^. Another potential unwanted outcome associated with anti-SARS-CoV-2 antibodies is severe acute lung injury mediated by IgG Fc-dependent macrophage recruitment in productively infected lungs, thereby skewing inflammation-resolving response ^22^. This risk of additional local inflammation should thus not be carried by pig IgG. Although unable to elicit ADCC, pig IgG are nevertheless endowed with a preserved complement-activation capacity. As enveloped viruses might be sensitive to direct complement-mediated lysis, administration of anti-SARS-CoV-2 pig antibodies might therefore lead to viral neutralization without targeting viral particles to inflammatory cells.

Our first attempt to use polyclonal glyco-humanized pig IgG against an emergent infectious disease was in an in vivo EBOLA virus model ^48^. Despite low levels of neutralizing antibodies had been obtained, we reported significant effect on the viral load and on animal survival demonstrating the high efficacy of the polyclonal approach. More recently, horse polyclonal IgG against SARS-CoV-2 spike RBD have been developed and demonstrated potent in vitro neutralizing capacity (Zylberman et al, Medicina (B Aires) 2020;80 Suppl 3:1-6). Horse IgG have to be used in their F(ab’)_2_ form to reduce the risk of serum sickness, i.e. without Fc fragments, a problem addressed in the current study by use of swine glyco-humanized antibodies, that cannot be recognized by anti-carbohydrate antibodies known to particularly evoke serum sickness, and also Fc-dependent macrophage activation ^22^. Yet using glyco-humanized swine IgG antibodies has the advantage over F(ab’)_2_ fragments to present an extended in vivo half-life comparable to that of other heterologous IgG immunoglobulins (unpublished data from our preclinical and clinical investigations).

We have provided proof-of-concept that pig hyperimmune glyco-humanized IgG against SARS-CoV-2 may be of clinical benefit and, in addition, that foreseen therapeutic doses will show efficacy at neutralizing virus while decreasing the level of evoked immune complexes and of serum sickness associated with the production of anti-Neu5Gc and anti-Gal antibodies. Data from previous experiments with CP transfer and from preclinical investigations in mice and primates ^37,49^ as well as preliminary data in the clinical arena ^35^ show that antibodies against RBD can indeed provide an immediate clinical benefit. Altogether, this information warrants clinical evaluation of the administration of pig glyco-humanized IgG against SARS-CoV-2 RBD in COVID-19 patients. Owing to the high antibody titers produced by CMAH/GGTA1 KO animals in a short period of time and to the significantly shorter, cheaper and easier pharmaceutical development of polyclonal antibodies as compared to mAbs or mAb cocktails ^50^, glyco-humanized anti-SARS-CoV-2 pig immunoglobulins have been among the first treatments specific to COVID-19 reaching the clinic (NCT04453384).

## Authorship

Conceived the study: OD, BV, JPS, EC.

Designed and supervised some experiments: OD, BV, CM, RB, SB, RTS, JMB, NG, MP, JPS

Performed the experiments: JR, GE, PJR, RD, CG, PD, ML, CM

Analyzed data: BV, JR, EL, JPS

## Acknowledgments

This work was supported by Xenothera and by grants from the Société d’Accélération et Transfert de Technologie Ouest Valorisation, French Region Pays de la Loire and Bpifrance.

---

**Supplementary Figure 1:**
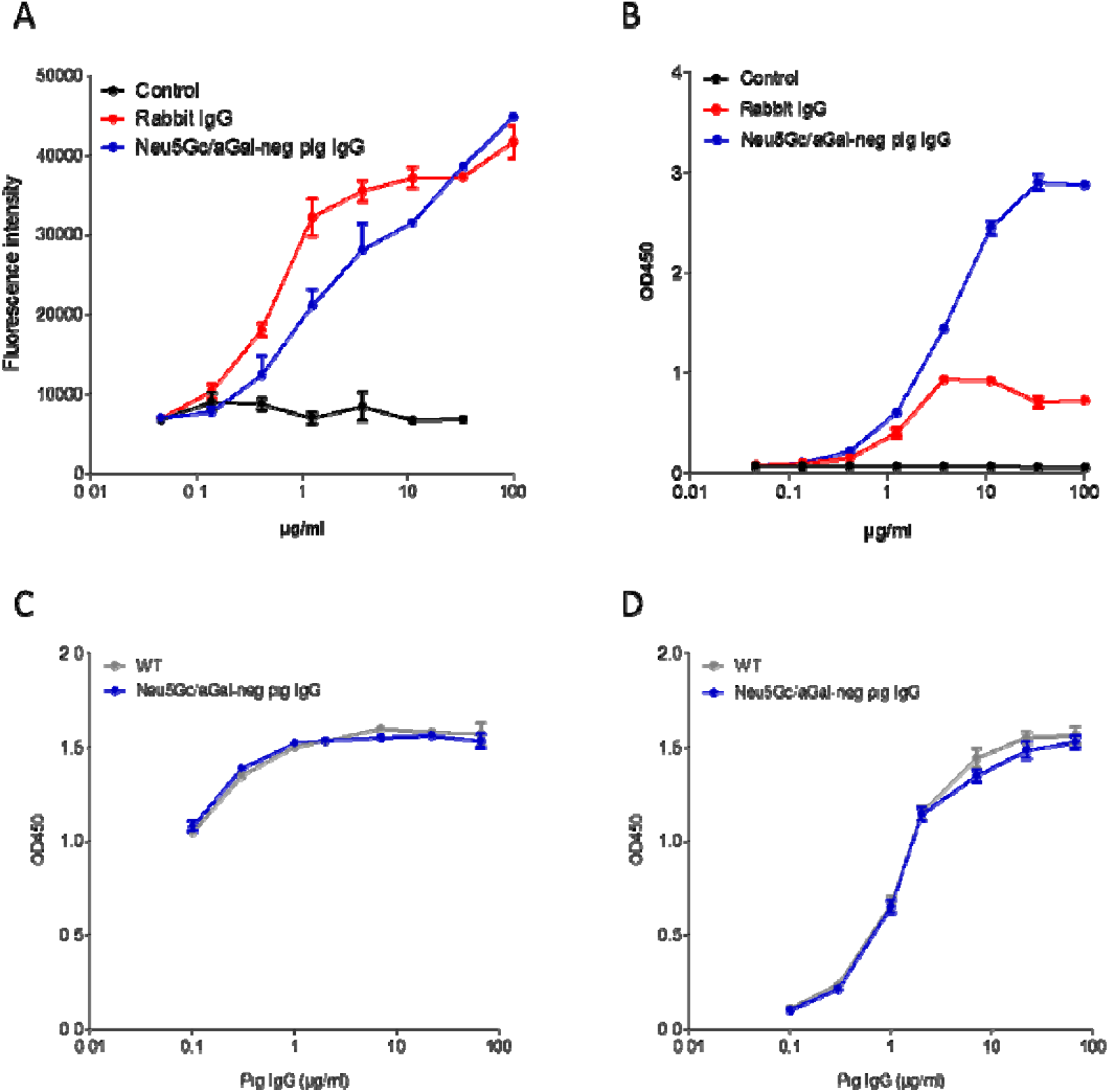
C1q binding of IgG from rabbit, WT pig and Neu5Gc/αGal-negative pig. A: Rabbit IgG or Neu5Gc/αGal-negative pig IgG were immobilized on plastic in ELISA plates. Equal coating intensity was controlled with a revelation using an Alexa fluor-conjugated Protein G and record of the fluorescence intensity. B: same coating as in A, with addition of human serum as a source of complement. Bound C1q molecules were detected with a sheep anti-human C1q and revealed with an HRP-labelled anti-sheep secondary antibody. C, D: WT and Neu5Gc/αGal-negative pig IgG preparations were immobilized on plastic at different concentrations. Similar coating of both IgG preparations was controlled by revelation with an HRP-labelled secondary antibody against pig IgG (C). Human serum as a source of complement was incubated and revealed as in B (D). Results are shown as means ± SD of triplicate experiments.

**Supplementary Figure 2:**
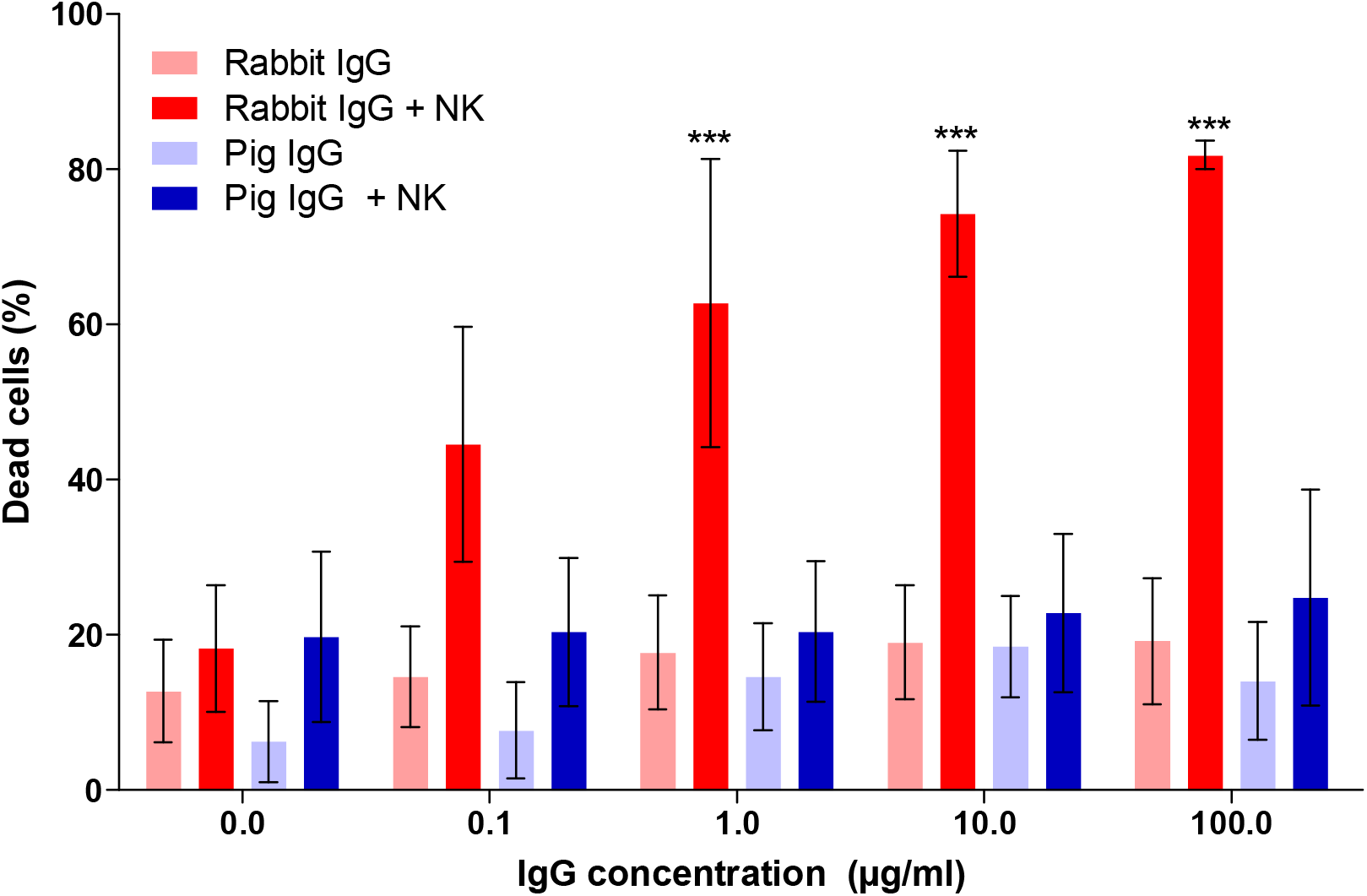
absence of ADCC activity of pig IgG towards human cell effectors. CMAH/GGTA1 KO pigs were immunized with human T cells. Immune Neu5Gc/αGal-negative IgG were purified and used in an ADCC assay with human NK effectors (effector to target ratio 10:1), in comparison with rabbit anti-human T cells IgG (Thymoglobulin®), at the indicated concentration. Results are shown as means ± SD of triplicate experiments. Two-way ANOVA followed by *post hoc* Bonferroni test. ***, p<0.01.

**Supplementary Figure 3:**
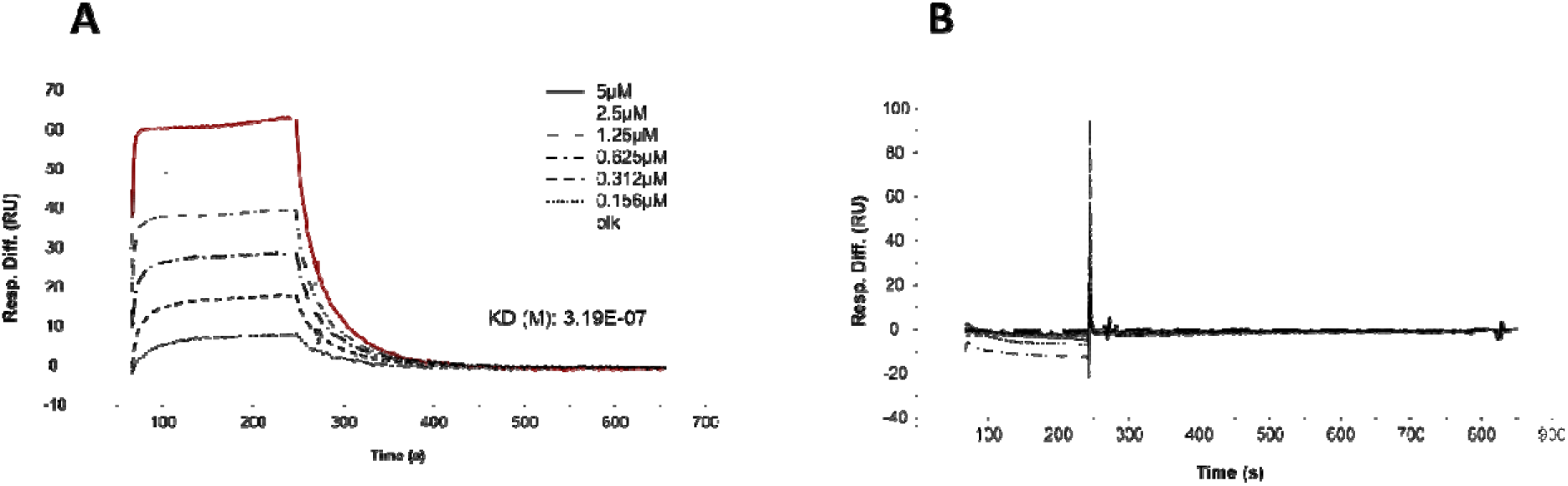
BIAcore surface plasmon resonance (SPR) analysis of pig and rabbit IgG binding to human Fc RIIIa (CD16) receptor. Sensorgram of rabbit (A) and pig (B) IgG interaction with human FcγRIIIa. Pig and rabbit IgG were immobilized on a CM5 sensorchip. Sensorgram were obtained by injections of different concentrations of human FcγRIIIa on the chip, with association and dissociation time intervals of 180s and 600s, respectively. Regeneration between cycles was performed by 100mM NaOH treatment for 45s. The resonance unit (R.U.) indicates the level of interaction. Dissociation constant (KD) for the rabbit IgG is indicated.

